# nzffdr: an R package to import, clean and update data from the New Zealand Freshwater Fish Database

**DOI:** 10.1101/2021.06.22.449519

**Authors:** Finnbar Lee, Nick Young

## Abstract

The New Zealand Freshwater Fish Database (NZFFD) is a repository of more than 155,000 records of freshwater fish observations from around New Zealand, maintained by the National Institute of Water and Atmospheric Research (NIWA). Records from the NZFFD can be downloaded using a web interface. The statistical computing language R is now widely used for data wrangling, analysis, and visualisation. Here, we present nzffdr, an open source R software package that: i) allows users to query and download data from the New Zealand Freshwater Fish Database directly in R, ii) provides functions to clean imported data, iii) facilitates the addition of information such as species names and Department of Conservation threat classification status, and iv) a workflow for visualising information from the NZFFD. The nzffdr package aims to standardise, simplify, and speed up a workflow likely already used in an *ad hoc* manner by scientists across New Zealand and abroad.

## Introduction

The New Zealand Freshwater Fish Database (NZFFD) contains over 155,000 observations of freshwater fish (plus freshwater shrimp and kōura) from across New Zealand dating back to 1901 (Crow, 2017). The observations typically include information on sampling location, date, time, fishing method, and the organisation that conducted the survey; less frequently information on the number and size of individuals caught is included. The database is a remarkable asset and is widely used to inform academic and governmental research and decision making (Goodman et al., 2014; Joy & Death, 2004). A limitation of the NZFFD is that it lacks some basic variables that individuals need to add each time they analyse NZFFD data; for example, species’ common and scientific names are not included (a 6 letter species code is included), nor is any other taxonomic information (e.g. family), threat classification status, and whether the species is native or introduced. Adding this information each time data are downloaded is not trivial and can be time consuming if many records are downloaded.

The statistical computing language R (R Core Team, 2020) has increased in popularity over the last decade, and is now one of the most common programming languages used by ecologists (Lai et al., 2019). R is typically used for data wrangling, analysis, and visualisation and is a popular tool for interrogating NZFFD data (Jellyman & Harding, 2012; Jones & Closs, 2015; Leathwick et al., 2006).

Here we present the nzffdr R package. We describe the features of each of the core functions in the nzffdr package and then illustrate how the functions can be used via an analysis of NZFFD data.

## Methodology

The nzffdr package has four core functions and four core datasets. The four core functions: i) import NZFFD data from in R. ii) clean up a variety of spelling inconsistencies and add a new variable “form” which describes the sampling habitat e.g. (river, stream, wetland etc.), iii) add missing information such as, family, genus and species names, common names, Department of Conservation threat classification status (Dunn et al., 2017) and whether the species is native or introduced and, iv) import and attaches associated REC data. The four built-in datasets are: i) a subset of 200 rows from the NZFFD that can be accessed without an internet connection and used for exploratory analysis, ii) the different fishing methods included in the NZFFD; it is possible to search the database using these terms so they are provided for reference, iii) scientific and common names of all species included in the NZFFD; the database can be searched by species name (using scientific or common names) so these are provided for reference, iv) a simplified version of the 1:150k NZ map outline available from Land Information New Zealand (https://data.linz.govt.nz/layer/50258-nz-coastlines-topo-150k/) to facilitate easy mapping of species’ distributions.

### Importing data: nzffdr_import()

The `nzffdr_import()` function is used to search the NZFFD and takes input arguments that align with the search options of the NZFFD web user-interface. There are seven search arguments:

1. `catchment`: this refers to the Catchment number, a 6-digit number unique to the reach of interest. Search using the individual reach number (e.g. `catchment` = “702.500”), or for all rivers in a catchment you can use the wildcard search term (e.g. `catchment` = “702%”).
2. `river`: search for a river by name; for example, to get all records for the Clutha (`river` = “Clutha”).
3. `Location`: search for river by sampling locality for example, to get all records from Awakino (`location` = “Awakino”).
4. `fish_method`: search by fishing method used, for example to get all records where fish were caught using a seine net (`fish_method` = “Other net - Seine”). There are currently 59 different possible options for fishing method, a list of all possible fishing method is available via the function *nzffd_method()*.
5. `species`: search for a particular species. There are currently 75 unique species in the NZFFD, a list of all possible species is available via the function `nzffd_species()`. Searches can be made using either common or scientific names and it is possible to search for multiple species at once. e.g. to search for Black mudfish use `species` = “Black mudfish” or `species` = “*Neochanna diversus*” and to search for Black mudfish and Bluegill bully use `species` = c(“Black mudfish”, “Bluegill bully”).
6. `starts`: starting search date.
7. `ends`: ending search date.

Not specifying the arguments will return all possible records. The `nzffdr_import()` function requires an internet connection to query NIWA’s database.

### Cleaning imported data: nzffd_clean()

While the data imported from NZFFD is generally does not have many errors there are some small inconsistencies (e.g. spelling of river and place names); the `nzffd_clean()` function aims to fix these errors. The first letter of all words in the columns “catchname” and “locality” are capitalised, and any non-alphanumeric characters are removed. Observations in the “time” column are converted to a standardised 24-hour format and nonsensical values (e.g. “0.677”) converted to “NA”. The organisation column (“org”) is converted to all lowercase and has non-alphanumeric characters removed. The NZMS260 map code (“map”) is converted to lower case and has any non-three-digit codes converted to “NA”. Observations in the catchment name column (“catchname”) are standardised, e.g. “Clutha River”, “Clutha r” and “Clutha river” all become “Clutha R”. Finally, a new variable “form” is added, which defines each observation as one of the following: creek, river, tributary, stream, lake, lagoon, pond, burn, race, dam, estuary, swamp, drain, canal, tarn, wetland, reservoir, brook, spring, gully or NA. The “form” variable is created by matching the above “forms” with the “locality” column; therefore, it reflects the description given by the “locality” variable.

### Filling in missing data: nzffd_fill()

Additional useful information can easily be added to the NZFFD dataset. The `nzffd_fill()` function adds columns giving the species’ common name (“common_name”), scientific name (genus + species, “sci_name”), “family”, “genus”, “species”, the Department of Conservation threat classification status (“threat_class”, [Dunn et al., 2017]) and whether the species is native or introduced (“native”). Additionally, if the “map” and “altitude” variables have some “NA” values; `nzffd_fill()` can fill most of these by extracting the relevant data from The NZMS260 map tiles (https://data.linz.govt.nz/layer/51579-nzms-260-map-sheets) and an 8m digital elevation model (https://data.linz.govt.nz/layer/51768-nz-8m-digital-elevation-model-2012) raster, respectively. This function requires an internet connection to query the 8m DEM.

### Adding River Environment Classification data: nzffd_add()

Finally, network topology and environmental information from the River Environment Classification (REC) database (Snelder & Biggs, 2004) can be added to the NZFFD data using `nzffd_add()`. This function takes the NZFFD “nzreach” variable and matches it against the corresponding “NZREACH” variable in the REC database, and imports all the associated REC data, adding 24 new columns to the NZFFD dataset. This function requires an internet connection to query the REC database.

### Illustration of nzffdr functionality

To demonstrate the utility of the nzffdr package we imported the entire NZFFD into R, cleaned up the imported data, filled in missing data, and added the REC database. We then highlight the usefulness of some of the new variables that nzffdr has added to the NZFFD dataset. Specifically, we map the distribution of native and introduced species, plot the relative proportion of records across habitat forms for each of the *Galaxias* species, highlighting their respective conservation status, and finally use the REC data to show the distance inland that each of the *Galaxias* species has been found.

All analysis was carried out using R v 4.1.0 (R Core Team, 2020). The package dplyr v 1.0.6 (Wickham et al., 2021) was used for data wrangling, ggplot2 v 3.3.3 (Wickham, 2016) for visualisation, and nzffdr v 1.0.0 (Lee & Young, 2021) used to access and tidy the NZFFD data. The code used to generate the results presented here is available via Figshare (https://doi.org/10.17608/k6.auckland.14776770.v1).

## Results and discussion

We plotted the distribution of introduced and native species records from the NZFFD (Fig. 1), where the introduced/native variable and the map of New Zealand are provided by the nzffdr R package. We then graphed the relative number of records occurring across 10 habitat forms for each of the *Galaxias* species, including information about each species’ threat classification status (Fig. 2). Habitat form, threat classification, and species common names have all been added to NZFFD data via the nzffdr package. Finally, distance to sea (km) at each of the locations of *Galaxias* species in the NZFFD have been observed at was plotted (Fig. 3). The distance to sea variable is added to the NZFFD data from the River Environment Classification database via the nzffdr package. This analysis illustrates some of the functionality offered by the nzffdr package.

**Figure 1.**
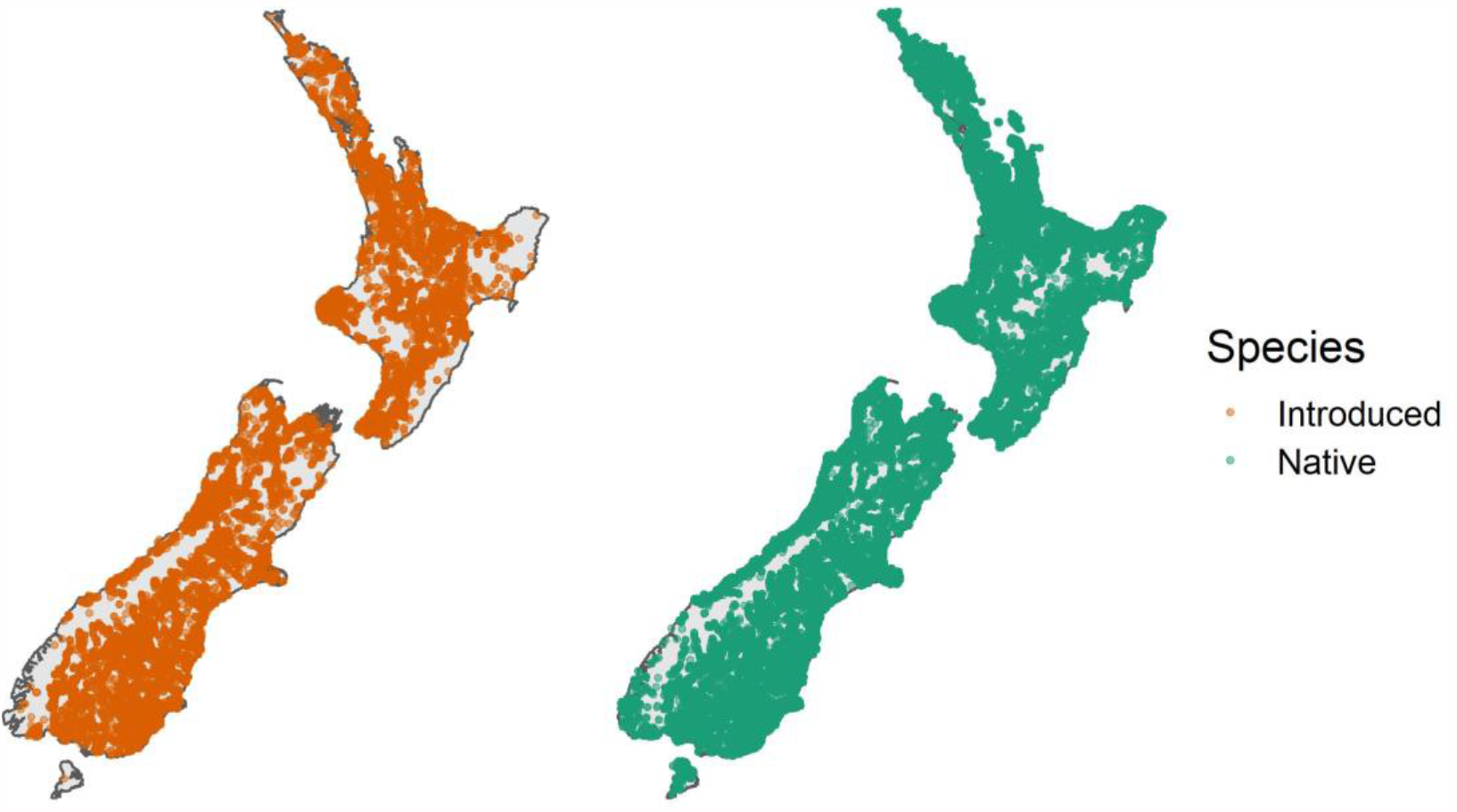
Distribution of introduced and native species records from the NZFFD, where the introduced/native variable and the map of New Zealand are provided by the nzffdr R package.

**Figure 2.**
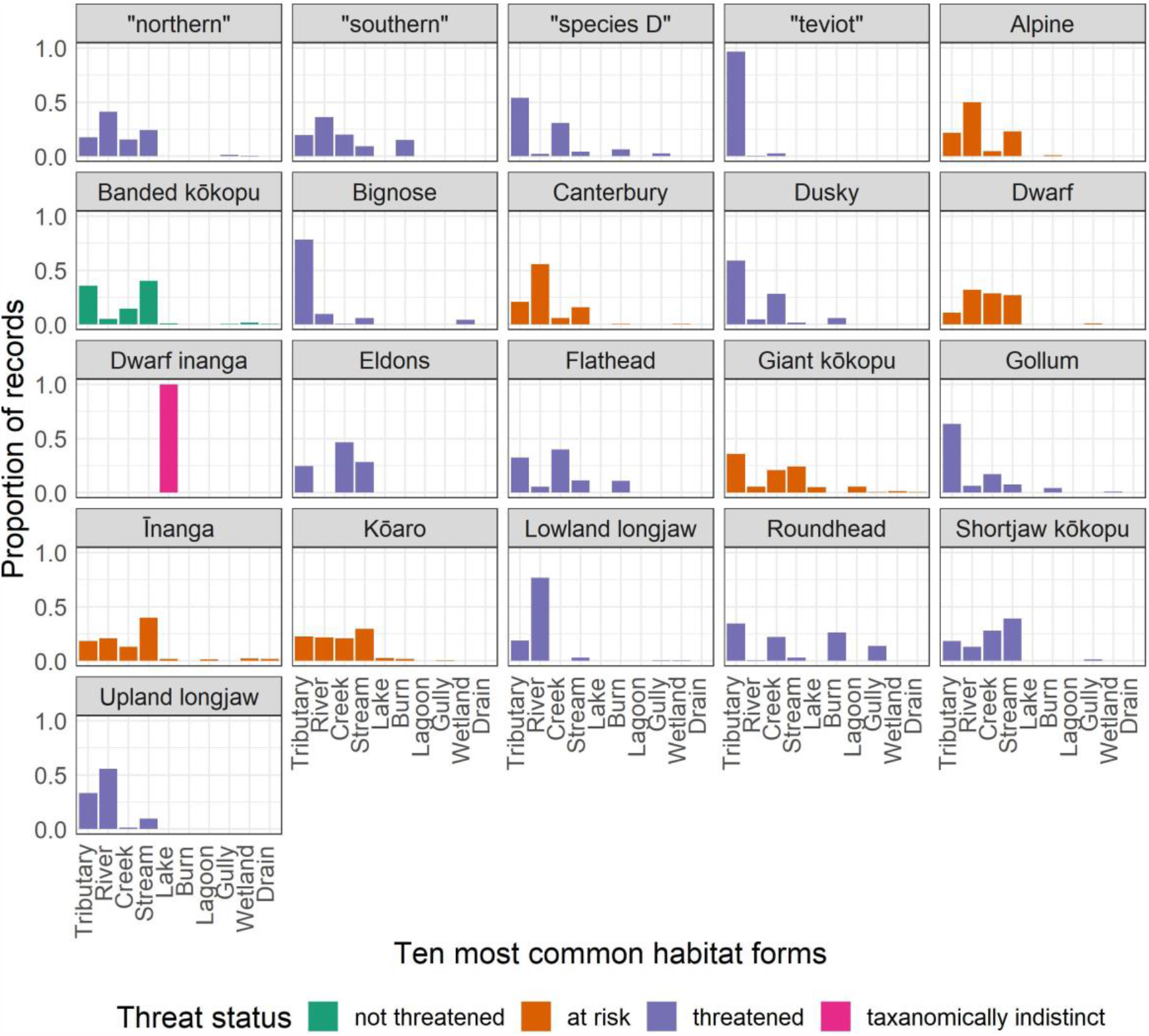
The relative number of records occurring across 10 habitat forms for each of the *Galaxias* species, the total number of observations for each species is given in parentheses. The habitat form and threat classification variables have been added to NZFFD data via the nzffdr R package.

**Figure 3.**
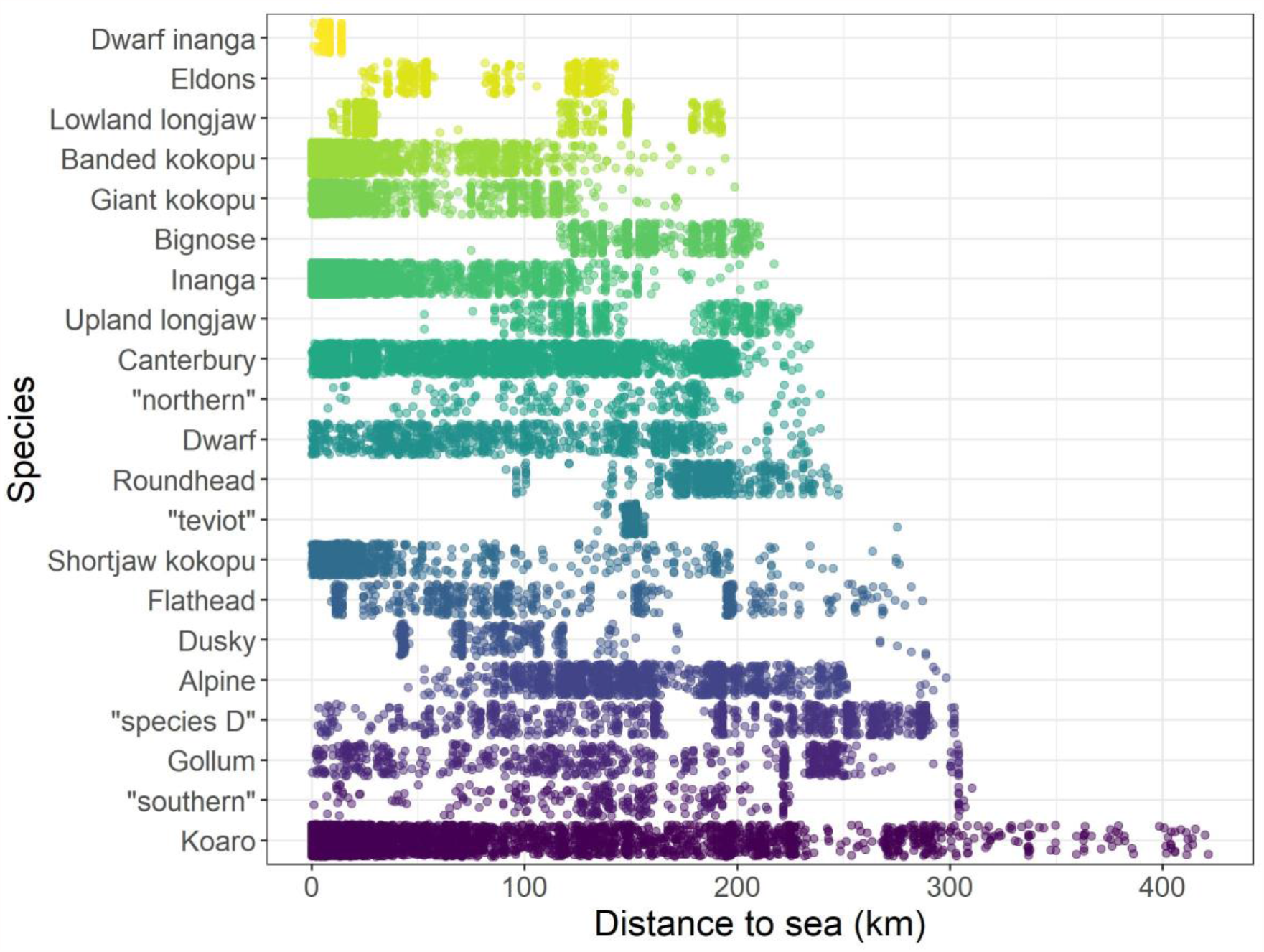
Distance to sea (km) that each of the *Galaxias* species in the NZFFD have been observed. The distance to sea variable is added to the NZFFD data from the River Environment Classification database via the nzffdr R package.

Here we have presented an overview of the nzffdr open source software package, which streamlines the importing, tidying, and adding of other important variables to the New Zealand Freshwater Fish database in R. This workflow is likely already being undertaken by researchers across New Zealand and overseas in an *ad hoc* manner. The nzffdr package speeds up this process and contributes to a reproducible workflow.

## Acknowledgements

The authors thank Quinn Asena, Jacqui Vanderhoorn, Craig Simpkins, Hayley Alena, George Perry, and André Bellvé for feedback on earlier versions of the code. This research was supported by use of the Nectar Research Cloud and by The University of Auckland. FL was supported by a George Mason Centre for the Natural Environment Research Fellowship.

## Data availability statement

The release version of the nzffdr software package described here is archived on the Comprehensive R Archive Network (https://cran.r-project.org) and the latest development version can be installed from https://github.com/flee598/nzffdr. The code used to produce the tables and figure in this manuscript is available via Figshare: https://doi.org/10.17608/k6.auckland.14776770.v1.

